# A *Trypanosoma brucei* ORFeome-based Gain-of-Function Library identifies genes that promote survival during melarsoprol treatment

**DOI:** 10.1101/2020.07.19.211375

**Authors:** M Carter, S Gomez, S Gritz, S Larson, E Silva-Herzog, HS Kim, D Schulz, GA Hovel-Miner

## Abstract

*Trypanosoma brucei* is an early branching protozoan parasite that causes human and animal African Trypanosomiasis. Forward genetics approaches are powerful tools for uncovering novel aspects of Trypanosomatid biology, pathogenesis, and therapeutic approaches against trypanosomiasis. Here we have generated a *T. brucei* cloned ORFeome consisting of over 90% of the targeted 7,245 genes and used it to make an inducible Gain-of-Function parasite library broadly applicable to large-scale forward genetic screens. We conducted a proof of principle genetic screen to identify genes whose expression promotes survival in melarsoprol, a critical drug of last resort. The 57 genes identified as overrepresented in melarsoprol survivor populations included the rate-limiting enzyme for the biosynthesis of an established drug target (trypanothione), validating the tool. In addition, novel genes associated with gene expression, flagellum localization, and mitochondrion localization were identified and a subset of those genes increased melarsoprol resistance upon overexpression in culture. These findings offer new insights into Trypanosomatid basic biology, implications for drugs targets, and direct or indirect drug resistance mechanisms. This study generated a *T. brucei* ORFeome and Gain-of-Function parasite library, demonstrated the libraries’ usefulness in forward genetic screening, and identified novel aspects of melarsoprol resistance that will be the subject of future investigations. These powerful genetic tools can be used to broadly advance Trypanosomatid research.

**IMPORTANCE:** Trypanosomatid parasites threaten the health of over 1 billion people worldwide. Because their genomes are highly diverged from well-established eukaryotes, conservation is not always useful in assigning gene functions. However, it is precisely among the Trypanosomatid-specific genes that ideal therapeutic targets might be found. Forward genetics approaches are an effective way to identify novel gene functions. We used an ORFeome approach to clone a large percentage of *Trypanosoma brucei* genes and generate a Gain-of-Function parasite library. This library was used in a genetic screen to identify genes that promote resistance to the clinically significant, yet highly toxic drug, melarsoprol. Hits arising from the screen demonstrated the library’s usefulness in identifying known pathways and uncovered novel aspects of resistance mediated by proteins localized to the flagellum and mitochondrion. The powerful new genetic tools generated herein are expected to promote advances in Trypanosomatid biology and therapeutic development in the years to come.

## INTRODUCTION

Trypanosomatids are a major parasitic lineage that include the African trypanosomes, American trypanosomes, and Leishmania spp. (family Trypanosomatidae, order Kinetoplastida), which collectively cause death and disease in millions of people living in tropical and sub-tropical regions(1). There are no vaccines against this family of parasites and the limited number of anti-Trypanosomatid drugs present ongoing challenges of host toxicity, complex treatment regimens, and burgeoning drug resistance(2).

Trypanosomatid parasites appear to have diverged from a shared ancestor around 100 million years ago. These evolutionarily ancient eukaryotes have highly divergent genomes from well-established model organisms with more than 50% of open reading frames (ORFs) annotated as hypothetical proteins(3). Of the 9,068 genes in the *Trypanosoma brucei* (African trypanosome) genome, 6158 are orthologous with both *Trypanosoma cruzi* (American trypanosome) and *Leishmania major*(3). While reverse genetics based on well-established models can promote discrete advances, forward genetics approaches have the potential to uncover important aspects of Trypanosomatid biology shared among orthologous genes.

*T. brucei*, the causative agent of Human African trypanosomiasis (HAT), has historically been the most genetically tractable of the Trypanosomatid parasites. For the past decade a whole-genome RNA interference (RNAi) knock-down library has been the primary forward-genetics tool in *T. brucei* resulting in identification of essential genes, genes associated with drug resistance and pathogenesis, and signaling factors critical to life cycle progression, to name a few(4–9). A strength of the RNAi library and associated RNA-Interference Targeted (RIT)-seq approaches is the identification of genes that result in a loss-of-function phenotype(10). A Gain-of-Function library approach may be more effective in the identification of genes whose expression is critical to the initiation of a cellular or developmental process. Gain-of-Function screens in other organisms have proved their usefulness in the identification of drug targets and resistance mechanisms(11–13). In basic biology they have been critical for discoveries in the areas of chromosome segregation and cell cycle, signal transduction, transcriptional regulation, cell polarity, and stem cell biology(14).

Traditional methods of overexpression library formation by cDNA synthesis and cloning are not viable for *T. brucei* as most gene expression regulation in Trypanosomatids occurs post-transcriptionally, with 5’ and 3’UTRs playing a major role in determining steady state levels of their associated transcripts(15). Existing *T. brucei* overexpression libraries generated by physical or enzymatic whole-genome fragmentation lack the ability to ensure complete ORF integration and can include unwanted regulatory elements(16, 17). ORFeome-based approaches, in which all ORFs in the genome are cloned for downstream applications, are powerful tools for the specific evaluation of gene effects whose proximal regulatory elements are excluded(18, 19). In addition, generation of an ORFeome can be applied to the downstream generation of multiple whole-genome methodologies, including yeast 2-hybrid libraries, tagging libraries, and inducible expression libraries for Gain-of-Function studies(20–23).

In this study we have taken an ORFeome-based approach to generate a *T. brucei* Gain-of-Function library for forward genetic screens. Melarsoprol was selected for a proof of principle genetic screen because of its clinical significance and unknown aspects of resistance and mode of cell killing. Melarsoprol, an arsenical compound, has long been used for the treatment of second stage (central nervous system) *T. brucei* infection(24). Second stage HAT infections caused by *T. b. gambiense* can now be treated by NECT (nifurtimox/eflornithine combination therapy) and the recently approved drug fexinidazole(2, 25). However, melarsoprol remains the only treatment for second stage *T. b. rhodesiense* infection, which rapidly progresses toward host death if left untreated. Melarsoprol treatment is burdened with high levels of host toxicity, challenging treatment regimens, and increasing reports of drug resistance and treatment failures(24). Melarsoprol is taken up into the cell by the P2 adenosine transporter (AT1) and aquaglyceroporin transporter (AQP2), which are mutated in most drug resistant isolates(24). Redox metabolism in Trypanosomatids is based predominantly on their unique dithiol molecule trypanothione and the trypanothione reductase(26). *In vivo*, melarsoprol is rapidly metabolized to trypanocidal metabolites including melarsen oxide, which binds trypanothione forming the stable adduct MelT(27), which is expected to have diverse effects on redox metabolism, ROS stress management, and the formation of dNTPs by ribonucleotide reductase(24, 26). Despite the established relationship between melarsoprol and trypanothione, the specific mode of melarsoprol cell killing remains unknown (24). Because the biosynthetic and redox utilization pathways contain enzymes unique to Trypanosomatids, they have been broadly explored as drug targets against American trypanosomes and Leishmania species(15, 28–32).

Here we present a description of the newly generated Gain-of-Function parasite library and describe its use in a screen for factors that increase parasite survival in the presence of melarsoprol. Library induction in the presence of melarsoprol resulted in the isolation of a specific survivor population consisting of 57 significantly overrepresented genes. Among these genes we identified the rate limiting enzyme of trypanothione biosynthesis (g-glutamylcysteine synthetase, *Tb927.10.12370*) whose established relationship with melarsoprol validates the Gain-of-Function libraries usefulness(33). In addition, we identified subsets of overrepresented genes encoding proteins associated with gene expression, the mitochondrion, and the flagellum whose association with melarsoprol had not been reported previously. Thus, the *T. brucei* ORFeome and resulting Gain-of-function library that we generated are now positioned to provide new insights into Trypanosomatid biology, pathogenesis, and drug resistance, which will promote the development of novel therapeutics.

## RESULTS

### Generation of a *Trypanosoma brucei* ORFeome

To generate a library consisting of all relevant ORFs from the *T. brucei* genome, start and stop sites for all *T. brucei* ORFs were obtained from available *TREU927* ribosomal profiling data for 9200 genes(34). The targeted *T. brucei* ORFeome filtered out 1956 ORFs that were unsuitable in size (<100 bp or > 4500 bp), an undesired product (ribosomal genes, *VSGs, ESAGS*, pseudogenes), or annotated as “hypothetical unlikely”. Known multidrug resistant channels (including MRPA whose overexpression causes melarsoprol resistance) were also excluded(33). PCR primers for the resulting 7245 targeted ORFs were designed in silico with *attB1* and *attB2* Gateway cloning sites with matched melting temperatures, synthesized, and resuspended in 21 separate 384-well plates that were organized by their anticipated ORF product size and gene annotations as either ‘known’ or ‘hypothetical’ (Table 1 and SUP. 1 – Oligo sequences).

**Table 1.** 384-well oligo plates.

| Plate name | Min ORF Length (bp) | Max ORF Length (bp) | # ORFs per plate |
| --- | --- | --- | --- |
| 0_hypothetical | 102 | 375 | 373 |
| 0_known | 147 | 591 | 315 |
| 1_hypothetical | 375 | 522 | 374 |
| 1_known | 594 | 849 | 365 |
| 2_hypothetical | 522 | 666 | 372 |
| 2_known | 849 | 1056 | 363 |
| 3_hypothetical | 669 | 822 | 382 |
| 3_known | 1056 | 1287 | 367 |
| 4_hypothetical | 822 | 993 | 372 |
| 4_known | 1287 | 1524 | 348 |
| 5_hypothetical | 993 | 1155 | 365 |
| 5_known | 1524 | 1857 | 342 |
| 6_hypothetical | 1158 | 1365 | 375 |
| 6_known | 1857 | 2337 | 362 |
| 7_hypothetical | 1368 | 1635 | 378 |
| 7_known | 2340 | 3504 | 376 |
| 8_hypothetical | 1635 | 1953 | 381 |
| 9_hypothetical | 1953 | 2508 | 381 |
| 10_hypothetical | 2508 | 3501 | 381 |
| hypothetical_last | 3504 | 4488 | 135 |
| known_last | 3507 | 4497 | 138 |
|  |  |  | 7245 |

Each ORF was PCR amplified in 384-well format from *LISTER427* genomic DNA. The general quality of the PCR reactions was assessed by the addition of SYBR green and measurement of the resulting Relative Fluorescence Units (RFU) (FIG. 1A). Based on the SYBR green assessment, initial PCR reactions resulted in the successful amplification of 94% of the ORFeome (6820/7245 ORFs) (FIG. 1B). To increase ORFeome coverage, we reamplified 429 failed PCR reactions and succeeded in producing 228 products resulting in a final total of 7039 PCR products amplified (97.2% of the targeted genes).

**Figure 1.**
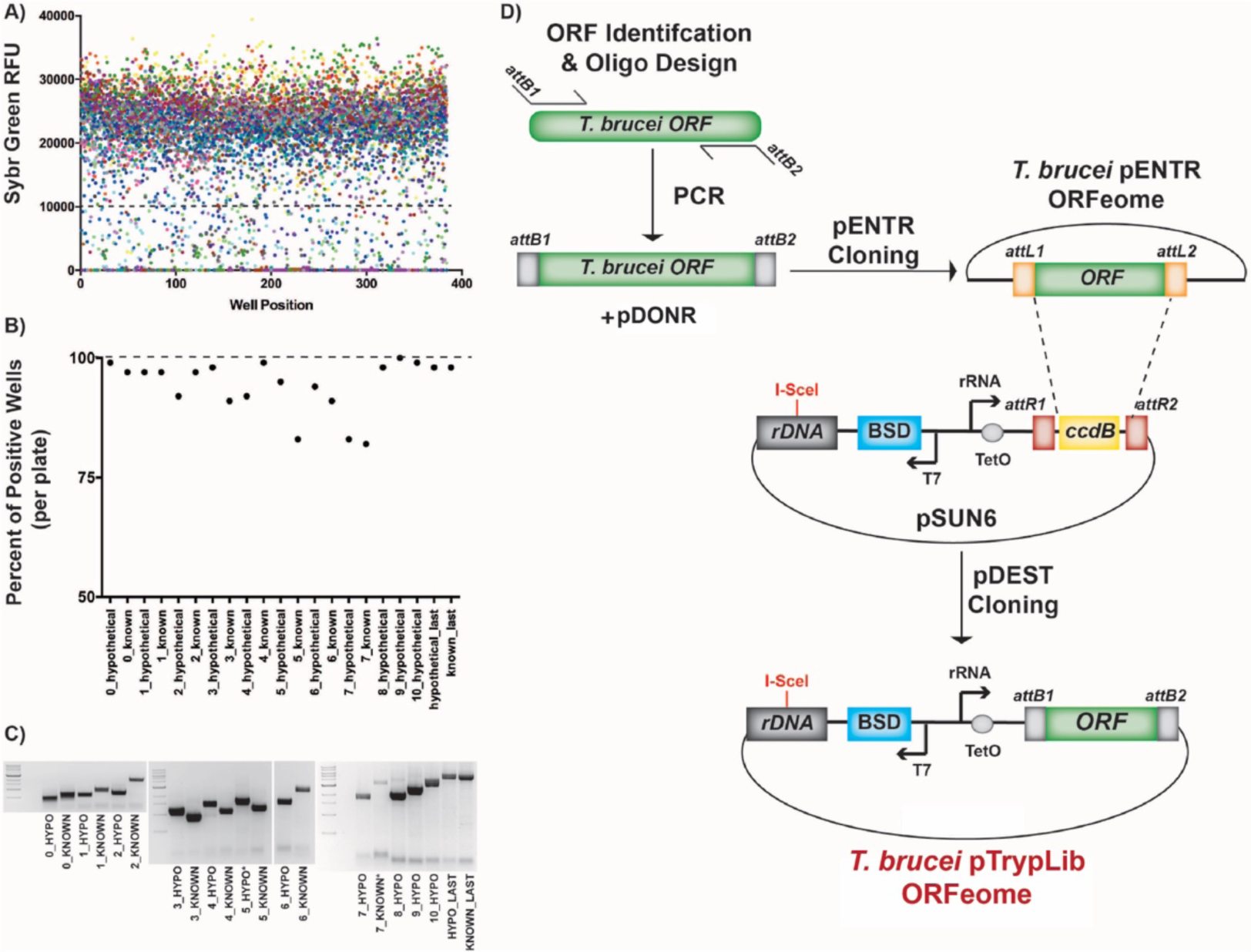
Generating a *T. brucei* ORFeome. A) Assessment of PCR amplification by SYBR Green Relative Fluorescence Units (RFU). Each color represents one of the 21, 384-well, plates and each dot represents a PCR reaction in a single well as measured by SYBR Relative Fluorescence Units (RFU). The graph is SYBR RFU vs. 384-well plate position. B) Percent of PCR positive wells (SYBR assessment) for each of the 21, 384-well plates, from the first time amplified. C) Agarose gel bands from each of the original 21, 384-well, PCR plates pooled prior to gel extraction and cloning, from the first time amplified, compared to 1 kb ladder DNA ladder. D) ORFeome cloning strategy: *attB* site addition to *T. brucei* ORFs during PCR amplification, BP Gateway cloning into pDONR221 to generate the pENTR ORFeome, and LR Gateway cloning into *T. brucei* specific pDEST (pSUN6, SUP. 2) to generate the complete pTrypLib ORFeome.

PCR reactions from each 384-well plate were pooled (10 µl from each well) into 21 corresponding PCR product pools, irrespective of the SYBR result, which maintained the product size range associated with each plate (Table 1). Each resulting size-sorted PCR pool was run on agarose gels and gel purified prior to Gateway cloning (FIG. 1C). Each size sorted pool of gel extracted PCR products were cloned into a standard pDONR Gateway cloning vector (pDONR221), as described (35), to generate the pENTR ORFeome library. The resulting pENTR libraries were then transferred into a *T. brucei* specific pDEST type vector with rDNA spacer targeting homology regions and a tetracycline-inducible system for ORF expression (FIG. 1D and SUP. 2 – pSUN6 map). The resulting library of ORFs cloned for *T. brucei* genomic integration was termed the pTrypLib ORFeome.

### Sequencing, assessment, and final coverage of the *T. brucei* ORFeome

The *T. brucei* pENTR and pTrypLib ORFeome harboring plasmids were each pooled and prepared for Illumina sequencing by tagmentation(36). In order to assess which of the 7245 targeted ORFs were not present in the pENTR and pTrypLib ORFeomes, we aligned the sequencing reads to the *TREU927* genome, removed PCR duplicates, and counted the number of reads corresponding to each targeted ORF. Because we knew that some of the targeted genes were highly similar or duplicated, we aligned the reads under two modes, one that required unique alignments and one that allowed multiple alignments. Both datasets were then assessed to determine how many genes were ‘missing’ from each library; defined as any targeted gene with zero aligned reads.

Initial analysis showed 1,845 missing ORFs from the pENTR library and 2,593 missing ORFs from pTrypLib (FIG. 2A - pENTR_1, pTrypLib_1, unique alignments). To increase the number of ORFs in the final library, PCR products corresponding to each missing ORF were isolated from the original PCR plates. The resulting eight additional size sorted ORF pools were gel purified, Gateway cloned (SUP. 3 – Table of cloning pools including ‘MISS_1-8’), sequenced by tagmentation, and analyzed as described above. The final ORFeomes were missing 457 from the pENTR library and 636 ORFs from pTrypLib (FIG. 2A – pENTR_Final, pTrypLib_Final, uniquely aligned reads, and SUP. 4 – Table of All Missing ORFs). The final pTrypLib ORFeome contains between 6,609 uniquely aligned and 6,803 multiply aligned *T. brucei* ORFs, resulting in 91-94% inclusion of the targeted ORFeome.

**Figure 2.**
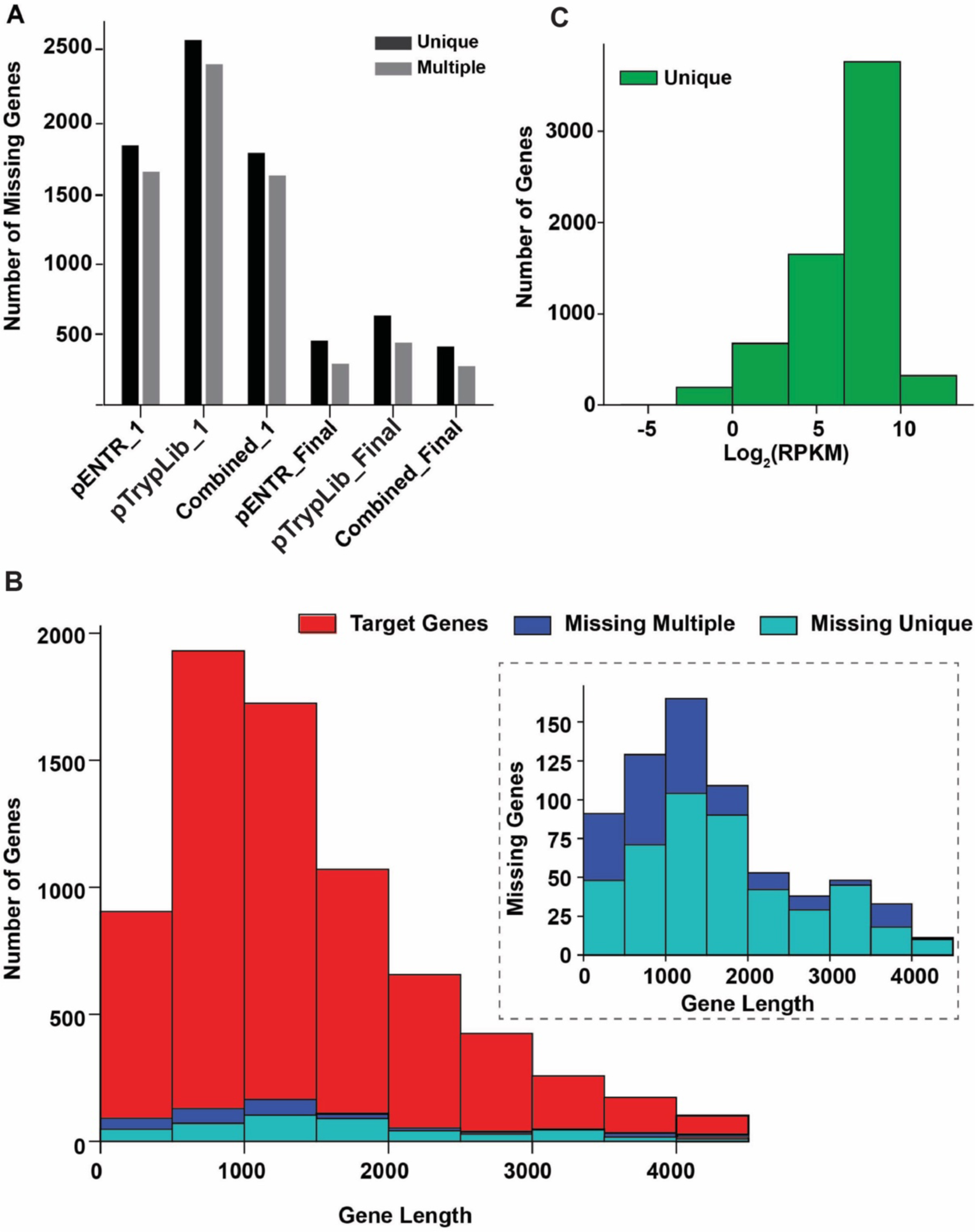
Assessment of pENTR and pTrypLib plasmid libraries. A) Bar graph showing the number of targeted ORFs with zero detectable aligned reads from the first round of cloning (pENTR_1 and pTrypLib_1) and after both rounds of cloning (pENTR_Final and pTrypLib_Final) using analysis generated from both uniquely and multiply aligned reads. Combined indicates a gene missing in both pENTR and pTrypLib. B) Histograms showing the distribution of ORF lengths for the target gene list (red) and the set of ORFs with zero detectable aligned reads after both rounds of cloning (labeled as missing). Analyses from unique (dark blue) and multiply (light blue) aligned reads are shown. Inset graph, target ORF lengths have been left out to better visualize the lengths of the missing ORFs. C) Histogram showing the distribution of normalized read counts for each ORF in the pooled pTrypLib plasmid libraries (Uniquely aligned reads shown, SUP. 5 both uniquely and multiply aligned reads).

To analyze whether large or small genes were overrepresented in the set of missing genes (unsuccessfully cloned ORFs), we compared the distribution of gene lengths between the target set of ORFs (red bars, FIG. 2B) and missing genes (blue and teal bars, FIG. 2B). The distribution of gene lengths was similar, indicating that cloning failure was likely independent of gene size.

Coverage of each ORF in pTrypLib was analyzed by count distribution based on number of reads aligned. Most ORFs resulted in log_2_(RPKM) values between 0 and 10 (FIG. 2C, unique, and SUP. 5 top panel, multiple). Thus, the number of poorly represented ORFs (RPKM < 1) is 195 for uniquely aligned reads and 369 for multiply aligned reads, representing 3% or 5% of all ORFs in the library, respectively. We then determined if ORF length affected representation in the library by plotting the log_2_(RPKM) value against ORF length (SUP. 5). No strong correlation was observed between ORF length and coverage in the pTrypLib ORFeome, with a best fit line showing a small negative slope for both unique and multiply aligned reads (−0.000673 and −0.000778, respectively). Thus, in general, shorter ORFs are not significantly more highly represented than longer ORFs, indicating that representation in the final libraries was not skewed by gene size (SUP. 5).

### A *T. brucei* Gain-of-Function parasite library

The pTrypLib ORFeome contains more than 6,500 tetracycline-inducible ORFs ready for *T. brucei* genomic integration at an rDNA spacer site. The landing pad (LP) system, developed for RIT-Seq library screens, was employed to ensure faithful integration into a single rDNA spacer site and high transfection efficiency (FIG. 3A), as described (6).

**Figure 3.**
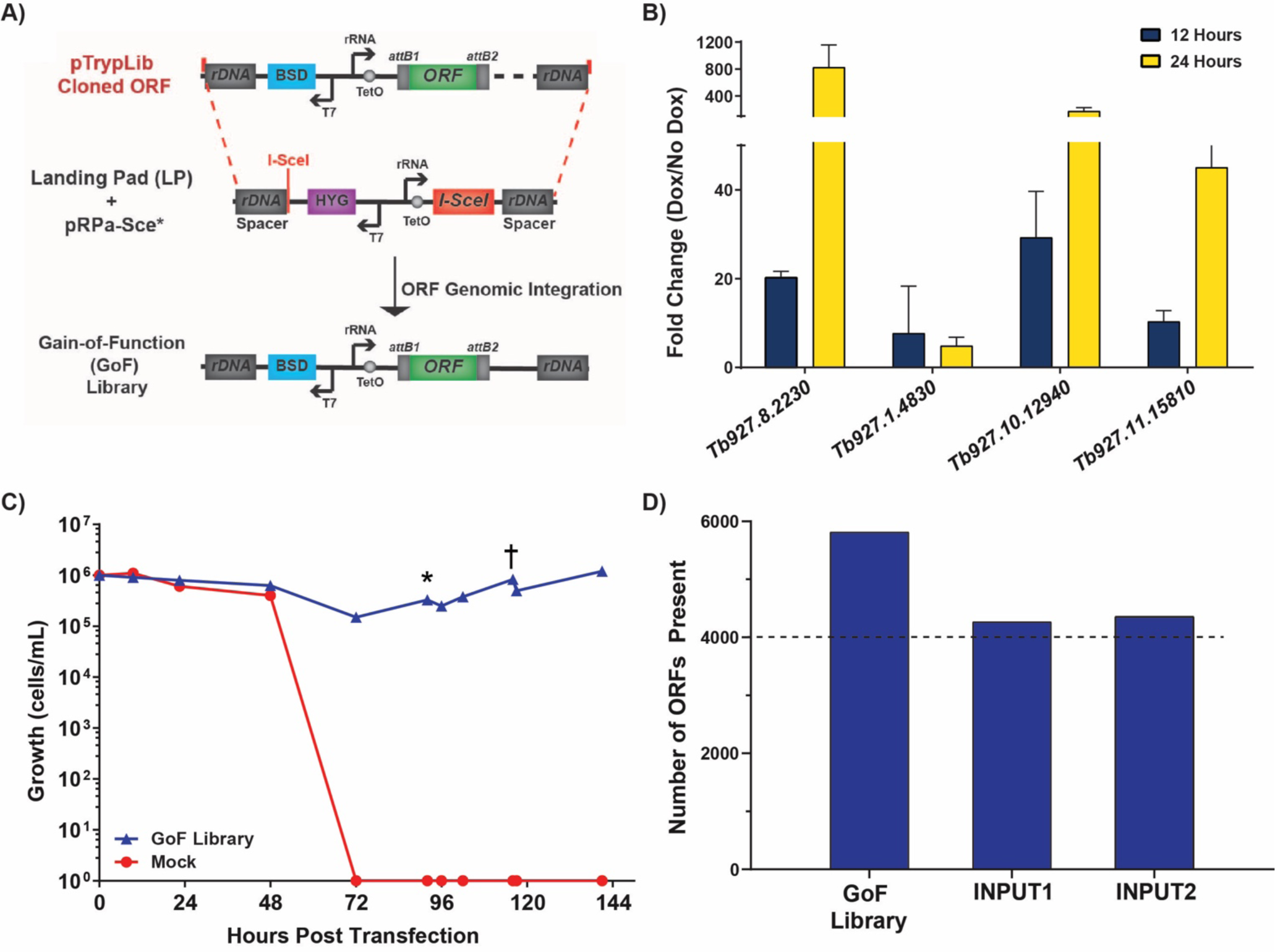
Generation and validation of the *T. brucei* GoF library. A) Transfection of pTrypLib ORFeome into parental Landing Pad (LP) cell line harboring pRPa-Sce* plasmid for I-SceI induced enzymatic cleavage of a single rDNA spacer site(6) B) Inducible expression of a Low Complexity GoF Library measured by RT-qPCR following 12 and 24 hours of doxycycline induction compared to uninduced cells (No Dox). C) Generation of the pTrypLib ORFeome based GoF parasite library. Graph shows the recovery of GoF library harboring cells (blue line) compared to mock transfection (red line) in blasticidin (BSD) added at Time 0, 12 hours post-transfection (* indicates cells spun and resuspended in 300 mL, † indicates addition of 500 mL). D) ent of the number of ORFeome genes present in the GoF library following nsfection (GoF library), in INPUT1 (following freeze thaw and minimal ion), and INPUT2, which used an alternative NGS protocol (See Methods).

Prior to transfection of the full pTrypLib ORFeome, we sought to verify inducible expression of this system using a low complexity library. The low complexity library was generated by transfecting a small number of equimolar pooled ORFs and recovered as a single population of parasites. Thus, we generated an ORF library with 1000 times less complexity than the complete pTrypLib. The low complexity library was then grown with or without doxycycline (Dox) induction for 12 or 24 hours prior to RNA extraction and RT-qPCR analysis to measure inducible expression of the transfected ORFs. ORFs showed increased transcript levels following Dox induction at 12 and 24 hours; 3 of the 4 ORFs analyzed resulted in approximately 10-30-fold increased transcript levels after 12 hours and 50-600-fold increase in transcript levels after 24 hours (FIG. 3B). Thus, the overall strategy of ORFeome exogenous transcription induction from pTrypLib cloned ORFs was deemed viable.

The full pTrypLib ORFeome was then used to generate an inducible *T. brucei* Gain-of-Function (GoF) library by transfecting 360 million cells (LP) and selecting with blasticidin (BSD)(6). 60 million cells survived transfection, which were then propagated to 3 billion cells over 3 days to generate the *T. brucei* GoF library (FIG. 3C – blue line). Illumina sequencing libraries are prepared using a custom P5 forward oligo containing *attB1* site complementarity and a universal P7 reverse oligo. Indexed products were Illumina sequenced using a custom oligo complementary to the *attB1* site (upstream of the introduced ORF. Thus, the resulting sequencing reads primarily correspond to the 5’ ends of the introduced ORF (SUP. 6A – Sequencing strategy). Immediately following transfection and recovery in blasticidin, the *T. brucei* GoF library consisted of 5,818 ORFs (FIG. 3D – GoF Library) and then approx. 4,300 ORFs following freeze thaw (FIG. 3D – INPUT1 and INPUT2). It is unclear if the apparent loss of approximately 1,500 ORFs arose through an artifact associated with a relatively low number of NGS reads returned from those samples or a true loss of content between library transfection and the subsequent thawing of frozen library.

### Isolation of melarsoprol survivors by Gain-of-Function genetic screening

We chose melarsoprol for a critical proof of principle GoF survivor screen because of its clinical significance, known interaction with trypanothione, and for the possibility of uncovering unknown aspects of its mode of action and resistance. To identify ORFs whose induced expression promote survival in the presence of lethal doses of melarsoprol, we tested three concentrations of drug on the LP parental cell line. Similar to previous reports, we observed that *T. brucei* LP cells died after 3 days in 35 nM, 5 days in 26 nM, and 7 days in 17 nM melarsoprol (17 nM is approximately two times the standard EC_50_ in culture and significantly less than concentrations used in clinical treatments) (FIG. 4A)(10). At 35 nM melarsoprol no survivor population emerged (FIG. 4B – Red dashed and dotted lines overlap). Thus, the 17 nM concentration was chosen to allow more time for induced ORF expression that might confer resistance.

**Figure 4.**
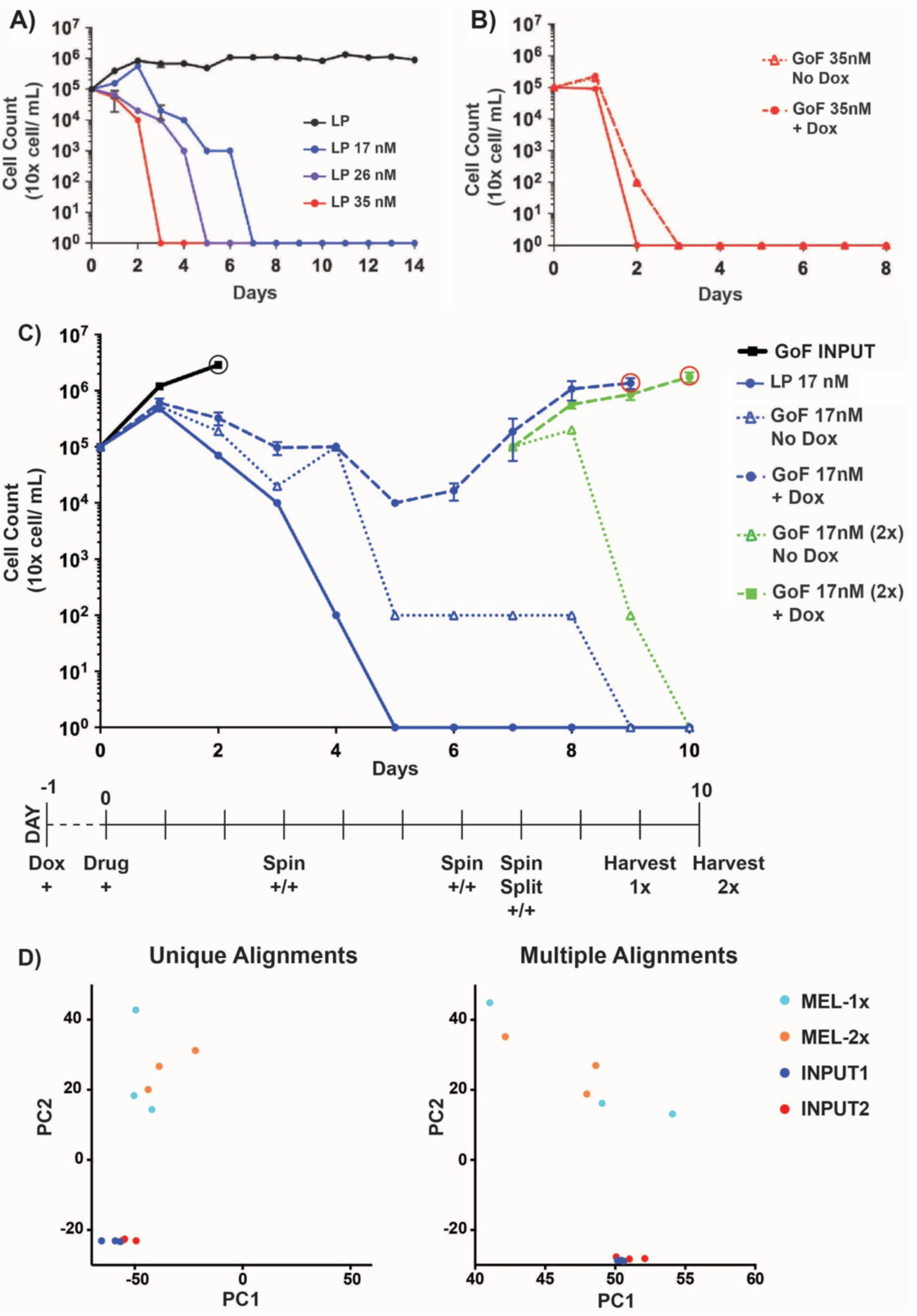
Isolation of melarsoprol survivor populations from a GoF screen. A) Growth of Landing Pad (LP) parental cell line in 17 nM (blue line), 26 nM (purple line), 35 nM (red line), or no (black line) melarsoprol. B) GoF library screen in 35 nM melarsoprol treatment: LP cell line (solid red line), uninduced GoF library (red dotted line), induced GoF library (red dashed line); dotted and dashed lines overlap. C) GoF genetic screen in 17 nM melarsoprol is shown. Timeline at the bottom of the graph indicates days on which either Dox (+ Dox), melarsoprol (+ Drug), or both (+/+) were added. All cultures (other than INPUT) were continuously grown in the presence of 17 nM melarsoprol. On days 3, 6, and 7 the triplicate cultures were centrifuged and resuspended in fresh media with melarsoprol and Dox, for induced, (noted as Spin +/+). Biological triplicate cultures are shown as: INPUT, untreated GoF library harboring cells grown for 3 days (black line), LP parental cell line (solid blue line), uninduced GoF library (No Dox, blue triangles on dotted line), induced GoF library (+ Dox, blue circles on dashed line), harvested on Day 9 (red circle on blue line) to produce MEL-1x. On day 7, biological triplicates from induced GoF library (+Dox, blue circles on dashed line) were split into two sets of triplicate samples, both in 17 nM melarsoprol, one of which was not further induced (No Dox, green triangles on dotted line). The other continued to be induced (+Dox, green squares on dashed green line) and was harvested on Day 10 to produce MEL-2x (red circle indicates harvest). D) Principle Component Analysis (PCA) comparing INPUT libraries (1 and 2) with libraries arising following continuous melarsoprol selection (MEL-1x and MEL-2x).

A GoF melarsoprol survivor screen was conducted for 10 days that included the following conditions: 1) Landing Pad parental cell line in 17nM melarsoprol, 2) untreated (no melarsoprol, No Dox) INPUT GoF Library, 3) uninduced (No Dox) GoF Library in 17nM melarsoprol, and 4) induced (+Dox) GoF Library in 17nM melarsoprol (FIG. 4C). Untreated GoF INPUT samples were grown in biological triplicates to Day 2, gDNA harvested, and represent the population of ORFs present before melarsoprol treatment (FIG. 4C – Black line). After 4 days of melarsoprol treatment the parental Landing Pad cell line had cell counts below the limit of detection (10,000 cells/mL) and from day 5 on showed no signs of life (FIG. 4C – Solid blue line). On Day 5, uninduced GoF Library counts were below the limit of detection, whereas induced GoF library resulted in a survivor population (FIG. 4C – Dotted blue line [No Dox] vs. Dashed blue line [+Dox]). The population of melarsoprol survivors arising from the induced GoF library began to replicate efficiently in the presence of drug following Day 5 and on Day 7 the triplicate samples were split into an additional 3 flasks that did not receive Dox induction (Green dotted line) and 3 with Dox added (Green dashed line); all continued to undergo melarsoprol treatment. Only Dox induced GoF library cultures were able to grow in the presence of melarsoprol (FIG. 4C - Blue and green dashed lines), suggesting that library induction promoted survival in these populations. The resulting populations of survivors, termed GoF ‘MEL-1x’ and GoF ‘MEL-2x’, were harvested for gDNA extraction at Day 9 or Day 10 (FIG. 4C – Red circles), respectively. Genomic DNAs from biological triplicate cultures of untreated INPUT (Day 2, black circle), GoF MEL-1x, and GoF MEL-2x cultures (9 cultures total grown to ∼1 million cells per mL 200 mL each) were prepared for NGS analysis.

We performed principle component analysis (PCA) on the resulting sequencing data using both unique and multiple alignments (FIG. 4D). The PCA analysis shows two clearly separated clusters for untreated and melarsoprol treated samples, with most biological replicates clustering proximal to one another. Melarsoprol treated samples were distinct from INPUT and show more variation between samples (FIG. 1D – INPUT1 and INPUT2 derived from the same source DNA, from which NGS libraries were generated using two elongation times. See Methods). We observed, at best, a weak negative association between gene length and normalized read count (slopes of −.00042 and −.00045 for unique and multiple alignment analysis, respectively), indicating that ORF representation in the library is largely independent of ORF length (SUP. 7)

### Identification of overrepresented Gain-of-Function ORFs in melarsoprol survivors

We reasoned that any gene whose induction contributed to melarsoprol resistance should be overrepresented in induced libraries generated from melarsoprol survivor populations. The designation ‘overrepresented’ indicates that the normalized read count for a particular ORF is higher in MEL-1x and MEL-2x than in INPUT samples, which suggests that overrepresented genes have enhanced survival in the presence of drug. To determine the fold change that represents a valid difference between melarsoprol treated and untreated conditions, we compared each of the three biological replicates of INPUT2 to one another and counted the number of ORFs with a 1.5, 2.0, or 4.0-fold change in normalized read count (FIG. 5A – INPUT2 replicates compared pairwise). By evaluating the biological variation between similarly treated replicates we found that while many ORFs varied in normalized read count by greater than 1.5-fold between replicates (more than 300), very few ORFs varied by greater than 4-fold (FIG. 5A, similar results obtained from INPUT 1, data not shown). Thus, we used a 4-fold change in normalized read count between melarsoprol treated and untreated samples as the minimum threshold for identifying an ORF as overrepresented (a ‘hit’) in this study.

**Figure 5.**
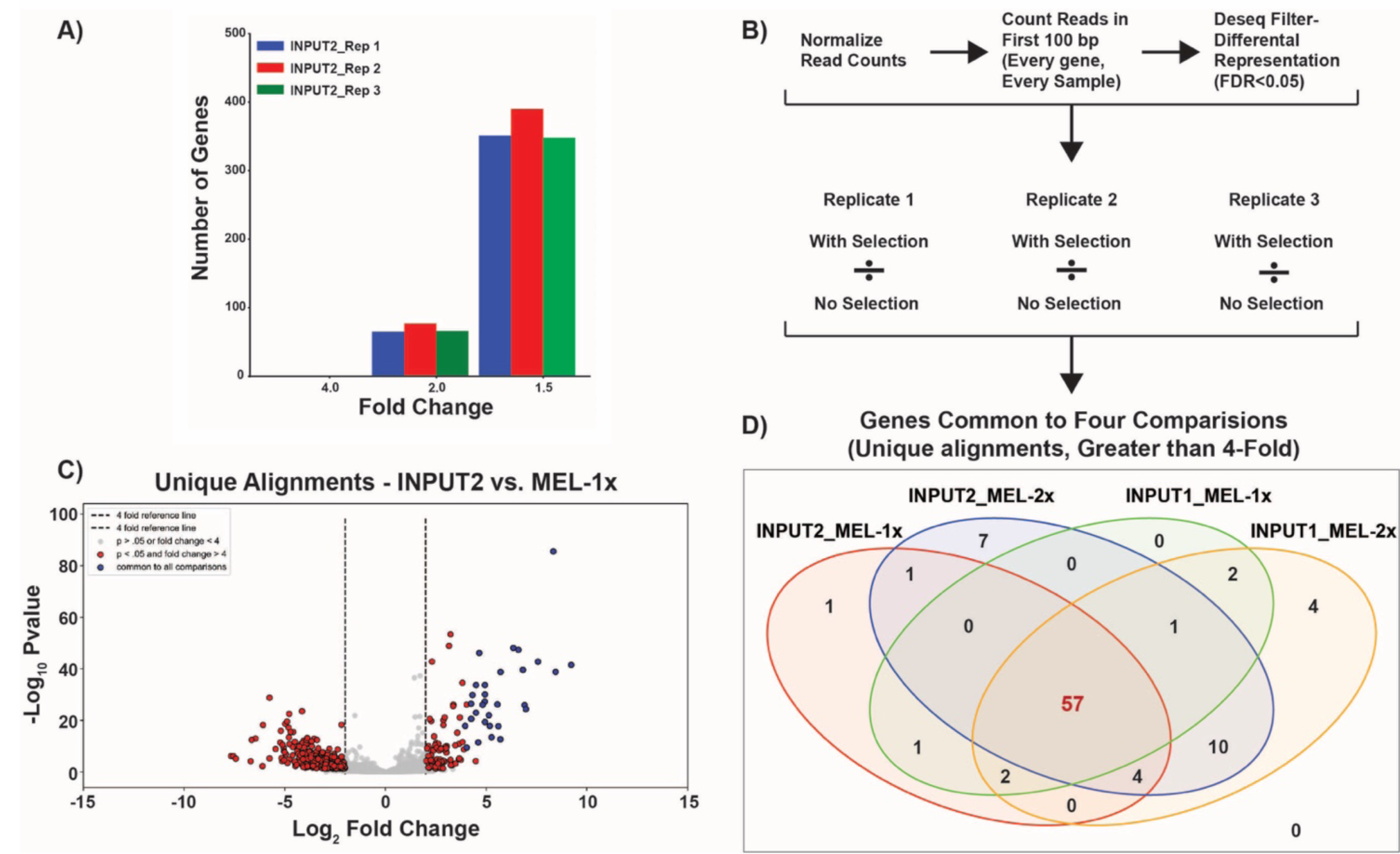
Identification of significantly overrepresented ORFs in melarsoprol GoF survivor populations. A) The number of genes with a fold change > 1.5, 2, or 4-fold is given for all three replicates of INPUT2: Replicate 1 (blue) vs. Replicate 2 (red), Replicate 2 (red) vs. Replicate 3 (green), and Replicate 1 (blue) vs. Replicate 3 (green). B) Hit calling pipeline to identify genes overrepresented in melarsoprol survivor populations. C) Volcano plot showing the −Log_10_(P adjusted value) vs. Log_2_ (Fold Change in normalized counts for the comparison of melarsoprol selected MEL-1x/INPUT2) for each ORF in the targeted library. Red dots represent fold changes >4 with a P adjusted significance of <0.05 and blue dots represent overrepresented ORFs common to all four comparisons described in panel D. D) Venn Diagram illustrates the significantly overrepresented genes common to each comparison between INPUT (INPUT 1 and 2) and melarsoprol treated (Mel-1x and Mel-2x) and shared among all comparisons between replicates, resulting in 57 overrepresented hits identified in melarsoprol survivor populations compared to input populations.

In order to identify ORFs overrepresented in melarsoprol survivor populations, we aligned reads from each of the 12 samples (3 reps of each: INPUT1, INPUT2, MEL-1x, and MEL-2x) to the genome, and then calculated the number of reads that fell within the first 100bp of each of the ORFeome targeted ORFs (SUP. 6B – Example read alignments). DESeq was used to normalize read counts between libraries and identify genes that were ‘differentially expressed’ between melarsoprol selected samples and INPUT samples with a p adjusted value of < 0.05 (FIG. 5B)(37). In this context, ‘differential expression’ simply refers to an ORF being overrepresented or underrepresented in the melarsoprol selected samples. To be considered as a potential hit, we required a minimum normalized read count of 5 in INPUT samples. To identify putative hits, we then asked which of the differentially represented ORFs with a p adjusted value < 0.05 had a normalized read count > 4-fold higher in 3 biological replicates of melarsoprol treated populations (MEL-1x and MEL-2x) compared to untreated populations (INPUT1 and INPUT2). Four different comparisons were analyzed using this pipeline: INPUT1 vs. Mel-1x, INPUT1 vs. Mel-2x, INPUT2 vs. Mel-1x, and INPUT2 vs. Mel-2x (FIG. 5B). Figure 5C shows a volcano plot of DESeq generated significance values versus fold change for the comparison between INPUT2 and MEL-1x. After hits had been called for each individual comparison, we identified the hits common between all 4 comparisons for both uniquely and multiply aligned reads (FIG. 5D and SUP. 8 – Tables of all comparisons). These analyses resulted in the identification of 57 overrepresented ORFs (uniquely aligned) in the GoF melarsoprol survivor populations compared to INPUT populations. In the comparison of INPUT2 vs. MEL-1x depicted in the volcano plot, we observe that these 57 ORFs common to all the comparisons were among the most highly overrepresented genes and with some of the lowest P adjusted values determined by DESeq (FIG. 5C – blue dots). Similar results were obtained for all comparisons between melarsoprol selected and input samples. We conclude that our analysis is reasonably robust at identifying the most consistent, statistically significant, highly overrepresented genes in melarsoprol treated samples compared to untreated samples.

### Melarsoprol resistance resulting from GoF hit overexpression

The 57 genes overrepresented in melarsoprol survivor populations are predominantly annotated as conserved hypothetical proteins or have putative functional assignments. Only two proteins among the melarsoprol GoF screen hits have known functions (*Tb927.10.12370* and *Tb927.11.15280*) though the functions of some annotated as putative are fairly well established (Table 2). To categorize all 57 genes, we utilized microscopic and proteomic localization data (curated through TriTrypDB) and available publications that addressed putative protein functionality. Based on this analysis we organized the hits into specific categories and found that the top three groups were associated with gene expression (16 genes), the mitochondrion (10 genes), and the flagellum (10 genes) (Table 2). The gene expression category could be further divided into proteins associated with splicing (5 genes), post-transcriptional regulation (5 genes), and translation (3 genes). It is important to note that categories based on localization were predominantly derived from data generated in insect stage (procyclic form) parasites, though some can also be confirmed from specific bloodstream form data (38–40). Based on these categories and the fold overrepresentation of each ORF in melarsoprol survivors we selected a subset of genes to analyze their effects on melarsoprol resistance.

**Table 2.**
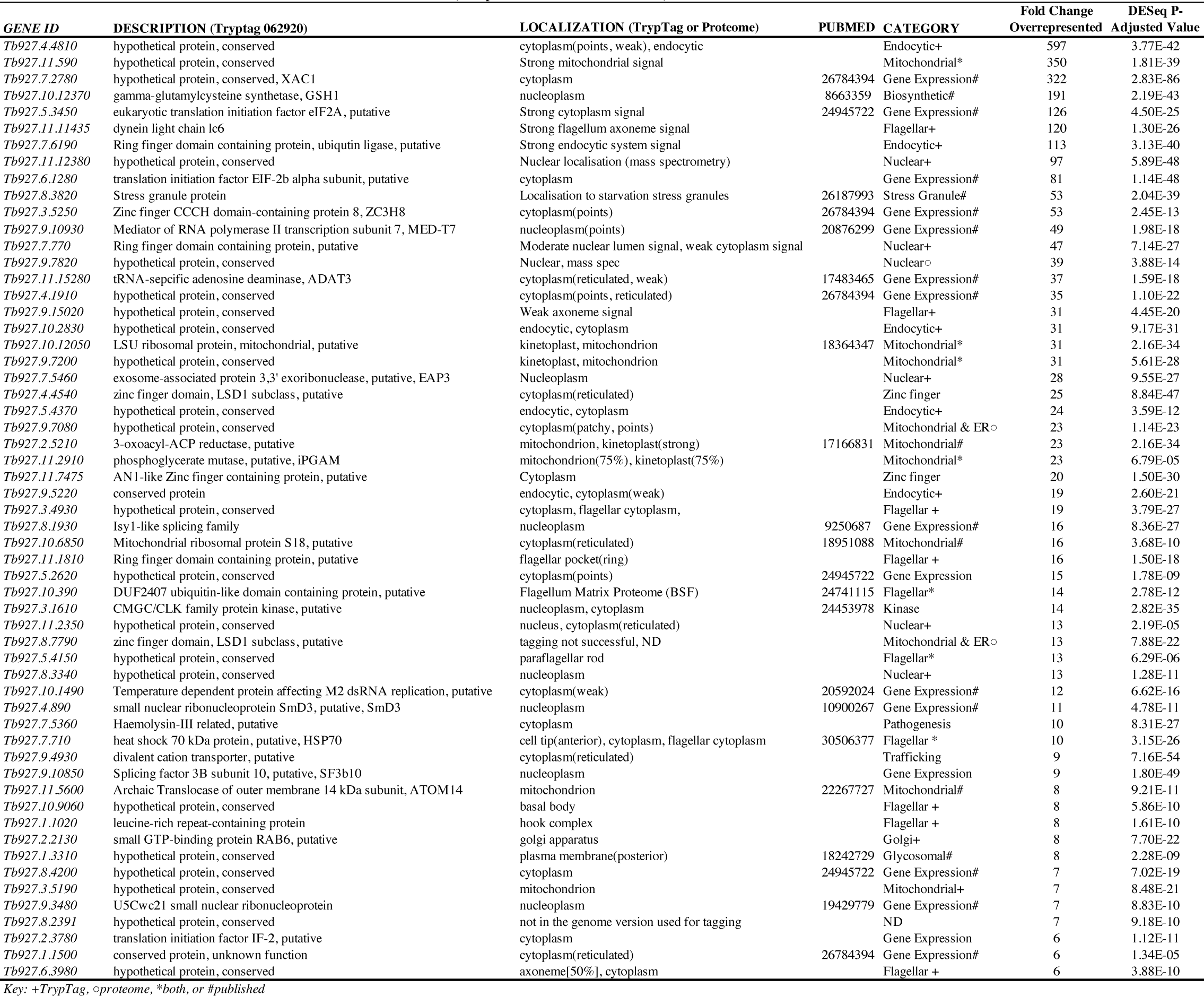
Hits overrepresented in melarsoprol survivors (Comparison of INPUT2 vs. MEL-1x)

We cloned a subset of overrepresented hits into a standard overexpression vector, transfected bloodstream form *T. brucei*, and analyzed their effects on melarsoprol resistance in cell viability assays (FIG. 6). The essential gene encoding g-glutamylcysteine synthetase (GSH1, *Tb927.10.12370*)(41), which is the rate-limiting step of trypanothione biosynthesis(28, 32, 33, 42), was 191-fold overrepresented in melarsoprol survivors (TABLE 2). Overexpression of GSH1 in *T. brucei* and other Trypanosomatids increases the concentration of intracellular trypanothione and has been shown to promote melarsoprol resistance under laboratory conditions(33). Our finding further supports the role of trypanothione as a primary intracellular target of melarsoprol and gives credence to the usefulness of the GoF library in the direct identification of drug targets. In our hands, overexpression of GSH1 resulted in an approximately 1.5-fold increase in the relative EC_50_ of melarsoprol (FIG. 6A and 6E).

**Figure 6.**
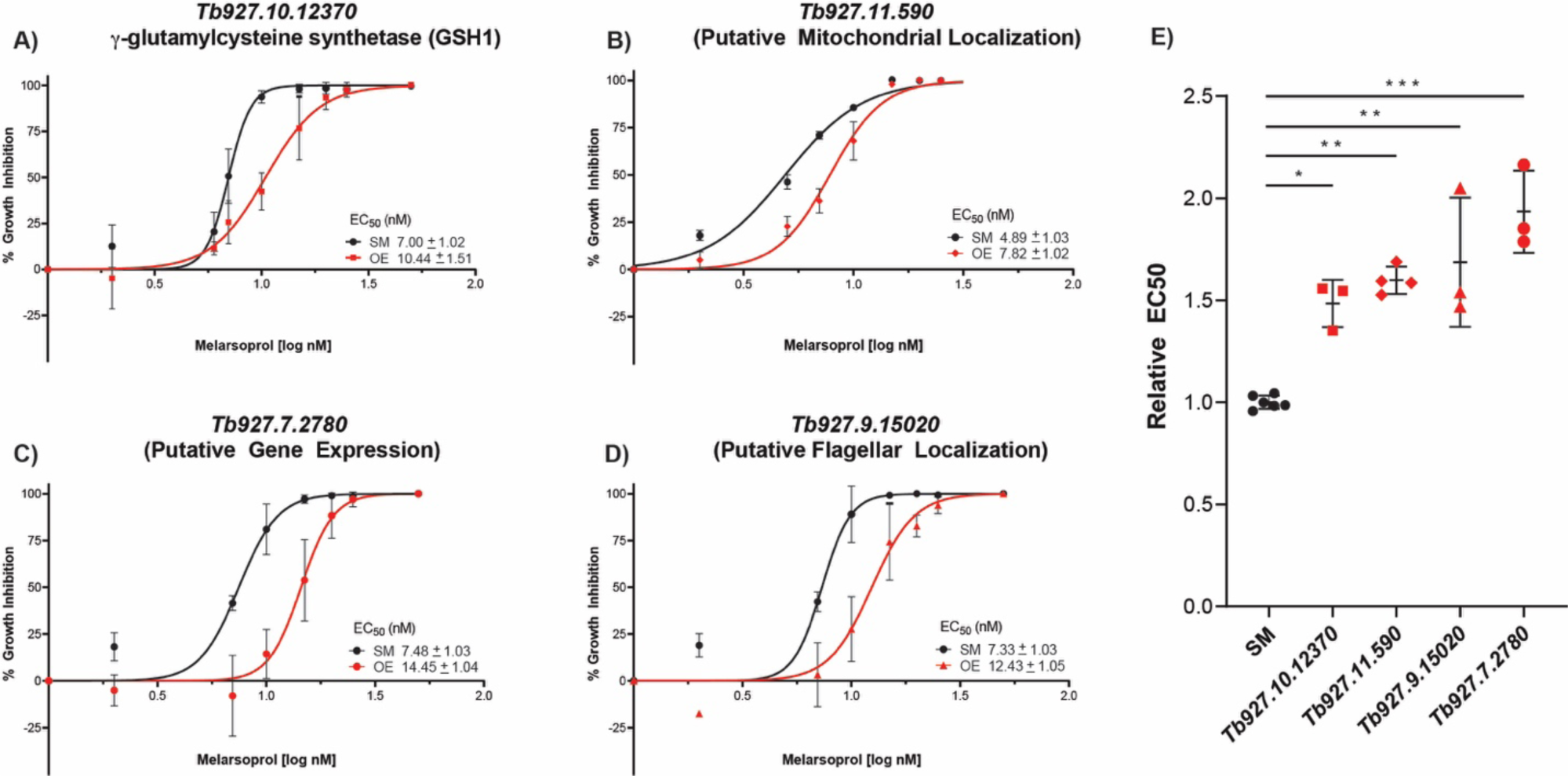
Melarsoprol resistance following gene overexpression. Induced expression of four hits (Red lines), in comparison with parental cells (SM, black line), during melarsoprol treatment with cell viability measured by alamarBlue assay to measure the resulting EC_50_: A) *Tb927.10.12370* (squares). B) *Tb927.11.590* (diamonds). C) *Tb927.7.2780* (circles). D) *Tb927.9.15020* (triangles). E) Relative EC_50_ following overexpression of each gene overexpressed gene for at least 3 biological replicates. P values were derived from one-way Anova analysis with Dennett’s multiple comparison test (* P<0.05, ** P<0.01, *** P<0.001).

We then evaluated the overexpression of three genes not previously linked to melarsoprol resistance, which were categorized as: mitochondrial (*Tb927.11.590*, 350-fold overrepresented), gene expression (*Tb927.7.2780*, 322-fold overrepresented), and flagellar (*Tb927.9.15020*, 31-fold overrepresented). The most pronounced effect was a 2-fold increase in relative EC_50_ of melarsoprol following the overexpression of *Tb927.7.2780*, which encodes the putative post-transcriptional activator XAC1(eXpression ACtivator 1) (FIG. 6C and 6E)(45). Overexpression of the mitochondrion localized protein encoded by *Tb927.7.2780* resulted in a greater than 1.5-fold shift in the relative EC_50_ of melarsoprol. Similarly, overexpression of the flagellar protein encoded by *Tb927.9.15020* resulted in an increase relative EC_50_ of melarsoprol (FIG. 6D and 6E). Together these results show that genes identified in melarsoprol GoF screening can promote drug resistance upon overexpression. Our results further support trypanothione as a major target of intracellular melarsoprol and implicate novel genes and mechanisms of melarsoprol resistance in Trypanosomatids.

## DISCUSSION

The forward genetics tools generated here address an urgent need to extend genomic functional characterization in *T. brucei* and its Trypanosomatid relatives. More than 30 years of genetic and biochemical studies in Trypanosomatids, 10 of which included the extensive use of an RNAi-based loss-of-function library, have produced key discoveries in parasitology and basic biology(46). Yet, with the functions of more than 50% of Trypanosomatid encoded genes largely unknown, many mysteries remain unsolved and more functional pathways must be delineated. Here we have generated two powerful tools for forward genetic approaches: an ORFeome consisting of over 6500 *T. brucei* ORFs and an inducible Gain-of-Function library harbored in *T. brucei* parasites, whose functionality was validated in a melarsoprol proof of principle screen.

Once in the cell, melarsoprol is metabolized into multiple forms, including melarsen oxide, which complicates the identification of drug targets and determination of its mode of cell killing. In this study we identified g-glutamylcysteine synthetase (GSH1, *Tb927.10.12370*) among our top hits, whose overexpression increases the intracellular concentration of trypanothione the primary intracellular target of melarsen oxide(33, 41, 47). Identification of GSH1 in the melarsoprol GoF screen adds additional support to this being a primary pathway in melarsoprol’s mode of action (FIG. 7 – T(SH)_2_ pathway). It is likely GSH1 overexpression generates sufficient levels of trypanothione to overcome melarsoprol inhibition (33). As the primary redox carrier in the cell, trypanothione functions in with the reduction of protein disulfides, detoxification of hydroperoxides, and the formation of dNTPs for DNA synthesis(26). Determining what aspects of trypanothione redox results melarsoprol-induced cell killing remains to be determined.

**Figure 7.**
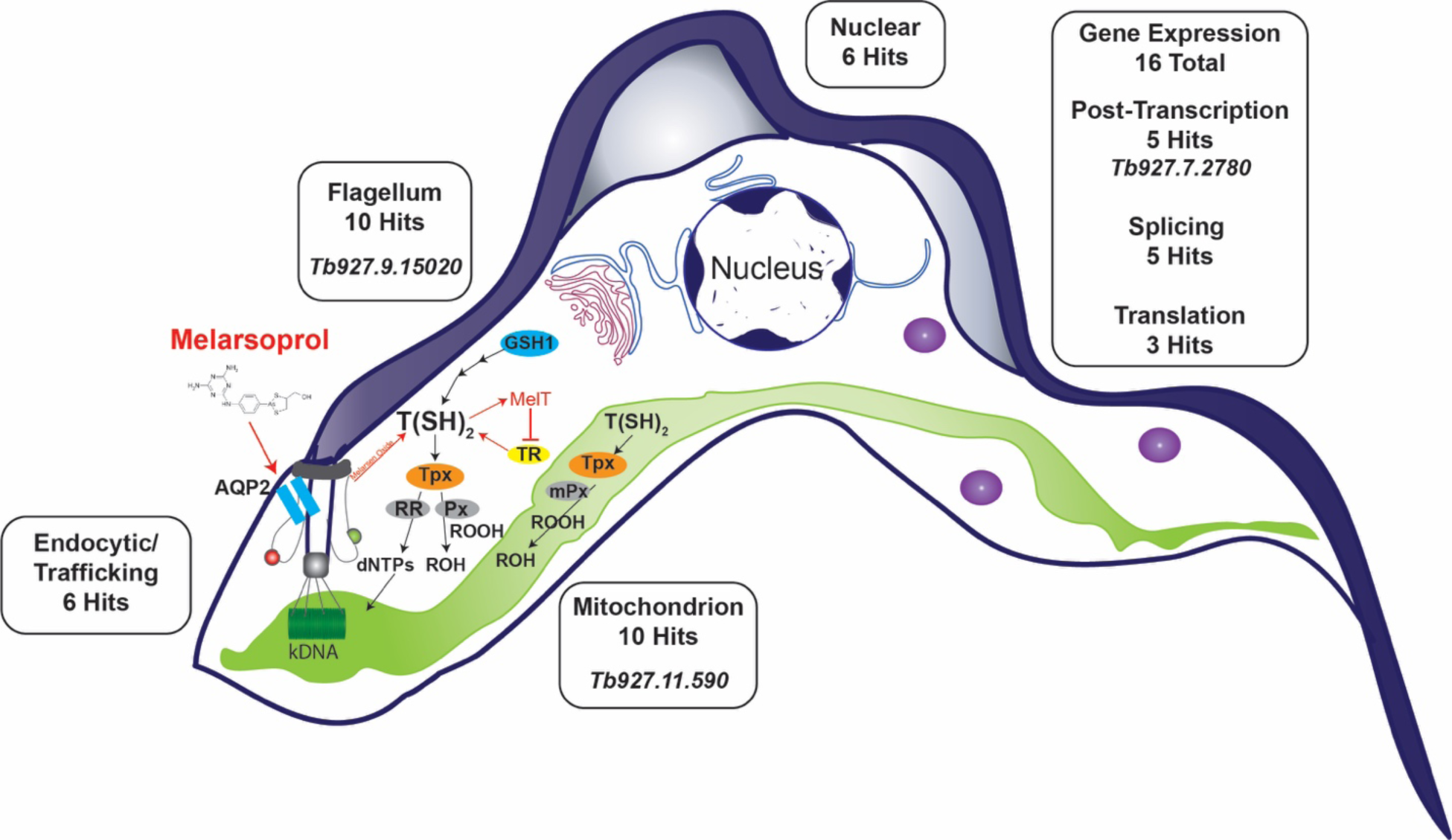
Categorization of GoF Hits. Hits arising from melarsoprol survival screening are shown proximal to bloodstream from *T. brucei* cell cartoon. Rectangular boxes indicate the number of hits occurring in each major GoF screen category (See TABLE 2 for details). Italicized gene names in boxes are shown for genes whose induced expression promoted melarsoprol resistance (FIG. 6). The cell diagram also highlights the flagellum and flagellar pocket with the melarsoprol transporter AQP2 localized as seen in bloodstream form (54, 55). Trypanothione (T[SH]_2_) biosynthesis and redox pathways are loosely depicted as follows: T(SH)_2_ biosynthesis is highly simplified showing the rate limiting enzyme GSH1, which was identified in the melarsoprol GoF screen; T(SH)_2_ provides reducing equivalents to tryparedoxin (Tpx), which is used to reduce disulfides (not shown), peroxidases (Px), and ribonucleotide reductase (RR) for the reduction of hydroperoxides and generation of dNTPs, respectively. T(SH)_2_ and Tpx are also utilized in the mitochondrion for redox reactions that include reduction of peroxidases (mPx). Melarsoprol uptake, conversion to melarsen oxide, binding with T(SH)_2_, to from the stable adduct MelT and its inhibition of trypanothione reductase (TR), which prevents the conversion of trypanothione disulfide back to T(SH)_2_ are all indicated in red. Green and red spheres at the flagellar pocket indicate import and export pathways, respectively.

Trypanothione biosynthesis and redox reactions primarily occur in the cytosol(26). Recently it was demonstrated that trypanothione and trypanothione reductase function in the mitochondrion (FIG. 7 – Mitochondrion in green), but these studies strongly suggested the requirement for unidentified oxidoreductases functioning in the organelle(48). Genes identified in the melarsoprol GoF screen suggest a previously uninvestigated connection between the drug and mitochondrion, though not entirely unanticipated based on trypanothione functions(26). The 10 melarsoprol GoF hits categorized as mitochondrial included β-ketoacyl-ACP-reductase (*Tb927.2.5210*, 23-fold overrepresented), which is required for fatty acid chain elongation in the mitochondrion as well as the production of the secondary redox carrier lipoic acid(49, 50). Here we have also shown that overexpression of *Tb927.11.590,* which encodes a mitochondrial protein with predicted oxidoreductase and catalytic domains, can increase the EC_50_ of melarsoprol (FIG. 6). It is intriguing to speculate that melarsoprol treatment may cause ROS or redox stress in the organelle, which might be alleviated by the overexpression of the mitochondrial proteins identified herein.

It is unclear at this time if overrepresented genes identified in melarsoprol survivors are direct targets of melarsoprol or they cause indirect effects that can promote resistance. Hits categorized as gene expression represent a complex list including genes associated with splicing, post-transcriptional activation and repression. The gene encoding XAC1 was among the top hits and has been reported to bind and stabilize mRNA transcripts to increase their translation (45). Overexpression of XAC1 increased the EC_50_ of melarsoprol (FIG. 6). We hypothesize that XAC1 overexpression may exert an indirect effect on melarsoprol resistance through the stabilization of transcripts that might be directly associated with the drug’s mode of action, perhaps those associated with trypanothione biosynthesis or utilization. It is important to consider the possibility that aspects of gene regulation might be directly targeted by metabolized forms of melarsoprol. The large number of putative splicing proteins (5 total) might suggest a drug interaction with the splicing complex that can be compensated for by protein overexpression(51–53).

The AQP2 transporter of melarsoprol and pentamidine is localized to the flagellar pocket in bloodstream form parasites (FIG. 7 – Blue rectangles)(54, 55). The large number of proteins localizing to the flagellum (10 genes) identified in melarsoprol survivors present the intriguing possibility that they function in aspects of drug transport. For example, overexpression of accessory proteins may result in reduced drug uptake that promotes resistance. Adding to this possibility, we identified 6 genes that were loosely categorized as endocytic or associated with intracellular trafficking. It would be useful to determine if any of these proteins affect the transport of trypanocidal drugs in a manner that might contribute to resistance. Flagellum proteins, mitochondrial proteins, and other categories of hits identified here present new testable hypotheses for future investigations that will likely uncover novel Trypanosomatid biology, drug targets, and alternative mechanisms of drug resistance (FIG. 6).

The functionality of the *T. brucei* ORFeome can be extended to generate additional genetic tools, such as, yeast two-hybrid libraries, tagging libraries, and dominant negative genetic screening approaches(18, 21, 22, 56). Based on the conservation of orthologous gene clusters among kinetoplastida(3), we expect the ORFeome could be used in other Trypanosomatids to generate orthologous Gain-of-Function libraries and other tools. The vast majority of genes overrepresented in melarsoprol survivor populations (∼80%, see SUP. 9) are conserved among sequenced Trypanosomatid genomes. This supports the use of these tools to broadly expand our understanding of gene functions in this family of parasites. We see the GoF library as a powerful new tool that can complement existing RNAi knock-down approaches and expand our understanding of drug targets and pathways of resistance. The tools and discoveries arising from this study are expected to support broad advances in basic biology, pathogenesis, pathways of drug resistance, and the discovery of new therapeutic targets in Trypanosomatid parasites.

## MATERIALS AND METHODS

Methods for ORFeome generation and assessment, Gain-of-function library assessment, and bioinformatic analysis of melarsoprol survivor populations are located in Supplement 10.

### Gateway cloning and plasmids

The pENTR library was generated by cloning each size-sorted PCR product pool into pDONR 221 Gateway Entry vector according to manufacturer’s specifications (https://www.thermofisher.com/us/en/home/life-science/cloning/gateway-cloning.html) and transformed into ElectroMAX DH10B cells by electroporation(35). The resulting transformants were plated on large LB plates containing kanamycin, assessed for efficiency of transformation, and bacterial colonies isolated from plates into LB liquid, which were split for maxi preps of plasmid and storage at −80°C in glycerol stocks. A *T. brucei* specific pDEST Gateway vector, pSUN6 (SUP. 2), was generated by introducing a ccdB Gateway cassette into a pLEW type vector(57) for incorporation into the *T. brucei* genome based on rDNA spacer homology, blasticidin selection, and ORF transcription from an rRNA promoter repressed by two tetracycline operators. Pools of pENTR plasmids harboring size sorted ORF populations were combined with pSUN6 in LR clonase reactions and transformed ElectroMAX DH10B cells by electroporation. The resulting transformants were plated on large LB plates containing ampicillin, assessed for efficiency of transformation, then bacteria and DNA isolated as described for pENTR steps above. The resulting plasmid libraries of pENTR and pTrypLib ORFeome Gateway cloning steps were assessed by NGS, described below. Following the initial assessment of ‘missing’ ORFs from both pENTR and pTrypLib cloning libraries, ‘missing’ PCR products were isolated from original plates, using a Perkin-Emer Janus Automated Workstation, to generate 8 new pools of size sorted PCR reactions (SUP. 3), which underwent the same series of Gateway cloning reactions described above and subjected to NGS analysis. The final NGS validated (below) pTrypLib library plasmids were pooled to generate a single pTrypLib ORFeome for introduction into the *T. brucei* genome.

### *T. brucei* cell lines, transfections, and GoF parasite library generation

Cell lines were generated from *Lister427* bloodstream-form trypanosomes derived from the ‘single marker’ (SM) line(58) and maintained in HMI-9 medium(59) under appropriate drug selection when indicated. A landing pad (LP) cell line was generated using plasmids gifted to us by the Alsford Lab and validated for inducible gene expression, prior to transfection with pRPaSce* as described(6, 60). LP parasites harboring the I-SceI cut site and I-SceI endonuclease gene targeted at an rDNA spacer were doxycycline induced to permit I-SceI cutting prior to pTrypLib ORFeome transfection by AMAXA Nucleofector(61). To generate the *T. brucei* GoF library described here, four 100ml flasks grown to ∼1 million cells/mL were AMAXA transfected with 10 µg pTrypLib DNA in four separate transfection reactions, which were then pooled into a single cell population in 500 mL of HMI-9 and recovered in a large roller flask, to which blasticidin was added 12 hours post transfection (FIG. 3C). An additional four transfections were completed in parallel with TE (Mock) to compare outgrowth with GoF Library transfection. The resulting blasticidin recovered GoF Library population was expanded to an 800 mL culture at ∼1 million cells per mL and saved in aliquots of ∼25 million cells per vial for future genetic screens. Cells were also sampled prior to freezing for NGS analysis (‘GoF library’, described below) and after freeze thaw (INPUT1).

Single gene overexpression cell lines were generated by cloning ORFs of interest into pLEW100v5-BSD (https://www.addgene.org/27658/), which following validation were digested with NotI and transfected into SM by AMAXA.

### Quantitative PCR assessment of ORF induction

Individual cloned ORFs were selected randomly from pTrypLib colonies plated on LB originating from the pool ‘2_known’, ORFs confirmed by traditional DNA sequencing, and DNAs arising from 4 individual ORF harboring pTrypLib vectors were transfected into LP harboring pRPaSce* by AMAXA as described above. This generated a ‘Low Complexity Library’ following transfection and recovery, which was split into No Dox and +Dox conditions for 24 hours, RNA was extracted, and cDNA prepared by Superscript III (ThermoFisher #18080044), prior to qPCR analysis. Quantitative PCR data was produced on a Bio-Rad CFX96 Real-Time PCR Detection System with iTaq Universal SYBR Green Supermix (Bio-Rad, #1725121). A forward primer anneals to the *attb1* site (5’-GGGGACAAGTTTGTACAAAAAAGCAGGCT) and reverse primer were unique to each ORF: Tb927.8.2230 (PRIMER: 5’-CACGGTTTTTGCCCATTCGT), Tb927.1.4830 (PRIMER: 5’-ATTTTTGCCGAAGCGCTTGA), Tb927.10.12940 (PRIMER: 5’-CCGTGATTCCCTGTCGACAT), and Tb927.11.15810 (PRIMER: 5’-CACCACCCGATGTACGGTAG). Because the forward primer anneals to the attB1 site present only in the pTrypLib backbone, only those mRNAs arising from the exogenous ORFs integrated at the rDNA spacer, rather than the endogenous ORF, can be detected. Fold changes in transcripts level with Dox and without Dox were plotted (FIG. 3B).

### Melarsoprol GoF library screening

GoF library cells were seeded for each condition at 1×10^5^ cell/ml, induced with doxycycline (1µg/mL) for 24 hours (for induced cultures, +Dox FIG. 4), and grown in HMI-9 medium containing Dox (when appropriate) plus melarsoprol at 17 nM or 35 nM melarsoprol (BoC sciences, CAS 494-79-1). Melarsoprol stocks were diluted in DMSO and cultures treated for the indicated duration (Fig. 4C bottom; a time bar indicates time points of replenishment of melarsoprol and/or Dox and time points of sample harvest). GoF library harboring cells were thawed from a single starting vial of approximately 25 million cells, propagated for 3 days prior to Day -1 Dox induction, and on Day 0 were split into 100 mL biological triplicates for untreated GoF Input samples (INPUT, No Dox), uninduced (No Dox), and induced (+Dox). Two elongation times were employed during PCR enrichment, INPUT1 for 75 seconds and INPUT2 for 20 seconds, to determine if amplification time resulted in a sequencing bias. Sequencing data was obtained in biological triplicate from INPUT1 and INPUT2 libraries (no melarsoprol treatment) and the two sets of melarsoprol selected parasites (MEL-1x and MEL-2x). GoF library harboring cells were recovered from each replicate and condition (INPUT, GoF MEL-1x, and GoF MEL-2x), genomic DNA fragmented, and prepared for ORFeome specific Illumina sequencing (SUP. 6).

### EC50 determination by alamarBlue

For EC_50_ determination, induced and uninduced cells were plated across a melarsoprol dilution series and viability was assessed after 72 hours using alamarBlue (ThermoFisher) as previously described(62). All experiments were performed in biological triplicate.

## ACKNOWLEDGEMENTS

The authors would like to extend our sincere thanks to the individuals and consortiums whose support made this work possible. Dr. Marilyn Parsons (now of Seattle Children’s Hospital) provided prepublication access to the TREU927 ribosomal profiling data that provided the gene starts and stops for all ORFeome targeted ORFs. Drs. Christine Clayton and Esteban Erben (Heidelberg University) provided their essential insights into methods for successful ORFeome Gateway cloning. Data provided directly from the Tryptag.org consortium was critical in the categorization of hits arising from melarsoprol GoF screening. Similarly, Tritrypdb.org was an essential resource throughout all stages of the work described herein. Finally, we would like to thank Dr. F. Nina Papavasiliou, whose support and generosity have been invaluable.

## Supplementary Figures

**Supplement 1.** Table of oligo pairs from all 21, 384-well plates. The sequences of all 7245 oligo pairs used to generate the *T. brucei* ORFeome are included with respect to their plate “name” and well position in each plate.

**Supplement 2.** Map of *T. brucei* specific pDEST cloning vector, pSUN6. Critical features of plasmid map are indicated, including rDNA spacer homology, T7 terminators, Blasticidin resistance cassette, rRNA promoter, two tetracycline operators, attR site for LR recombination with pENTR library, and the 3’ GPEET and 5’ Aldolase (long) UTRs.

**Supplement 3**. Table of all final ORF pools used to make the pENTR ORFeome. Following the first pENTR_1 and pTrypLib _1 ORFeome assessments, ORFs missing from libraries were identified and reselected from original 21 PCR plates to form an additional 8, size sorted pools. The original 21 pools, 2 pools of PCR reamplifications (NEG_PICKS) and 8 ‘missing’ ORF pools (1-8_MISS) collectively result in 31 pools for pENTR cloning. The size ranges and total number of ORFs per pool are indicated.

**Supplement 4**. Tables of all missing genes from final pENTR and pTrypLib ORFeomes. Consists of four sub-tables for all the ORFs identified as missing from either pENTR or pTrypLib by analysis of uniquely or multiply aligned reads: MISSING_pENTR_UNIQUE, MISSING_pENTR_MULTIPLE, MISSING_ pTrypLib _UNIQUE, AND MISSING_ pTrypLib _MULTIPLE.

**Supplement 5.** Assessment of the pTrypLib ORFeome. A) Histograms showing the distribution of normalized read counts in the pTrypLib ORFeome in analyses using uniquely aligning reads (left) and allowing multiple alignments (right). B) The length of each ORF in the pTrypLib Orfeome was plotted against the normalized read count of the ORF. A best fit line was calculated using linear regression (shown in white). For uniquely aligned reads this was y = 7.623751 - 0.000673x and for multiply aligned reads this was y = 7.306181 -0.000778x.

**Supplement 6**. Sequencing strategy and results from transfected libraries. A) Library preps from pTrypLib transfected parasites proceed by first fragmenting genomic DNA, ligating Illumina adaptors, and amplifying library introduced ORFs using a primer complementary to the *attB1* Gateway cloning sequence and the standard Illumina barcoded reverse primer. Sequencing proceeds using a custom forward primer complementary to the *attB1* Gateway cloning sequence. B) Red rectangles in the first row represent annotated genes from a section of the *T. brucei* chromosome 5. Bars in subsequent rows represent reads that align to the genes in the first row. Most reads align to the first 100bp of the gene, as expected from the library prep and sequencing strategy.

**Supplement 7**. Assessment of coverage in the GoF INPUT library. A) Histograms showing the distribution of normalized read counts in the sequencing libraries from INPUT 1 in analyses using uniquely aligning reads (left) and allowing multiple alignments (right). B) The length of each ORF in the pTrypLib Orfeome was plotted against the normalized read count for the INPUT1 sequencing library. A best fit line was calculated using linear regression (shown in white). For uniquely aligned reads this was y = 6.046291 -0.000423x and for multiply aligned reads this was y = 6.083779 - 0.000452x.

**Supplement 8.** Tables of all data pertaining to genes overrepresented in melarsoprol survivor populations.

**Supplement 9.** Table of conserved orthologs among the 57 genes overrepresented in melarsoprol survivors in other Trypanosomatid.

**Supplement 10**. ORFeome generation, NGS, and Bioinformatics Methods

